# Assessment of variability in the human urinary nucleic acid adductome

**DOI:** 10.64898/2026.10.14.759185

**Authors:** Alexandra Keidel, Jazmine Virzi, Mina Mckee, Yuan-Jhe Chang, Mu-Rong Chao, Chiung-Wen Hu, Leo Stockfelt, Gerd Sallsten, Lars Barregard, Marcus S. Cooke

## Abstract

Assessment of the urinary nucleic acid adductome provides a valuable, non-invasive approach for evaluating whole-body exposure to genotoxins. To support its application to exposome research, a clear understanding of biological variability independent of exposure is required. In this study, we analyzed a urine biobank from 40 healthy, non-smoking adults. Samples were processed using dual-phase solid-phase extraction, which retains a broad spectrum of nucleic acid adducts, and analyzed using an Orbitrap IQ-X platform. Data processing was performed using MZmine, FeatureHunter, and MetaboAnalyst. Across approximately 160 adducts per sample, the number of recovered adducts remained consistent between collection times, indicating that qualitative characterization of the urinary nucleic acid adductome is largely unaffected by diurnal variation. Most adducts exhibited stable abundances across sampling times. Diurnal variation was observed only in males, with 10 adducts showing significant differences between collection times, while females showed no temporal differences. Of the total adducts detected, only 32 differed significantly by age and/or sex. Age-related differences were more pronounced, particularly in males, where all significantly altered adducts were present at higher intensities in the younger cohort (20–29 y) compared with the older cohort (30–60 y). These findings support the use of spot urine samples for adductomic studies, favoring first-void urine. Collectively, these findings provide the first comprehensive assessments of biological variability within the human urinary nucleic acid adductome and support its application as a robust biomonitoring tool for investigating endogenous and environmental exposures across diverse human populations.

## Introduction

The exposome represents the totality of exogenous and endogenous exposures that an individual encounters throughout their life.^1–3^ Collectively, these exposures influence genome integrity by inducing chemical modifications to DNA, RNA and (2’-deoxy)ribonucleotide pools.^2,4,5^ For instance, oxidative stress, inflammation, and xenobiotic metabolism can generate reactive intermediates that covalently modify nucleobases, forming nucleic acid adducts.^5–7^ Epidemiological studies have connected environmental and lifestyle exposures, such as tobacco smoking, consumption of red meat and pollution to elevated levels of adducts.^8–16^ If unrepaired, these adducts can induce mutations, chromosomal instability, and ultimately contribute to carcinogenesis.^17–24^ A wide range of cancers have been linked to lifestyle-associated exposures, and nucleic acid adductomics offers a means to characterize the mechanisms and molecular consequences of these risk factors.

To avoid the downstream consequences of DNA adduct persistence, repair pathways remove these lesions from genomic DNA, and the arising modified nucleobases or 2’-deoxyribonucleosides are excreted into urine.^25–30^ Although their origins are less well established, the presence of modified RNA products in urine imply that analogous processes occur for RNA. ^31^ Targeted analysis of individual DNA and RNA adducts in urine has been used previously to determine the whole body exposures to nucleic acid damaging agents and processes. ^31^ However, unlike targeted analyses of individual lesions, untargeted nucleic acid adductomics captures the broad spectrum of DNA and RNA products arising from multiple damage pathways, allowing assessment of the body burden of adducts and temporal patterns without prior assumptions about lesion identity. Thus, the urinary nucleic acid adductome serves as a highly valuable non-invasive biomarker comprehensively reflecting the internal dose of all exposures, and the effect on target molecules (DNA, RNA and NTPs). ^32^

The urinary nucleic acid adductome may exhibit circadian variation due to intrinsic biological rhythms/processes and/or external exposures; excluding the former allows for more reliable assessment of the latter. Intrinsic factors which may affect the urinary nucleic acid adductome include circadian regulation of the kidney,^33^ fluid intake, as well as gene expression and transporter activity in renal tissues, which fluctuate across the 24 h cycle, influencing urine volume, pH, and solute composition.^34,35^ Additionally, external exposures from the environment may vary over the day.

In practice, urinary nucleic acid adducts concentrations are most commonly measured using single “spot” samples rather than 24 h collections. Diuresis, defined as increased urine production following fluid intake, may reduce solute and adduct concentrations in daytime samples compared to overnight collections. To compensate for changes relative to urine volume, adduct concentrations are commonly adjusted using the urinary concentration of creatinine or specific gravity (SG). Normalization to creatinine is based on the assumption that its excretion is independent of urinary flow rate; however, this assumption is not quite valid.^36,37^ Instead, creatinine excretion rates (mmol/h) exhibit circadian variation, reflecting changes in urinary flow as well as contributions from muscle metabolism.^38,39^ These factors introduce potential bias when creatinine is used as a correction metric in time-resolved analyses. Despite these limitations, both creatinine and SG remain widely used normalization approaches, as they reduce variability associated with urine dilution and improve comparability of urinary adduct measurements across samples collected at different time points.^38,40^

Although the diurnal variation of the urinary nucleic acid adductome has not been reported, variation in individual nucleic acid adducts has been previously studied. Notably, urinary levels of 8-oxo-7,8-dihydro-2′-deoxyguanosine (8-oxodG), a oxidatively generated DNA lesion, have been reported to range 8.4 to 32.88 nM in a population of 15 healthy volunteers.^41^ Studies of 24 h urine collections indicate that total daily excretion of 8-oxodG is relatively stable and spot samples correlate with the 24 h gold standard.^40^ Variability reflects differences in urine dilution and excretion dynamics across the day. Accordingly, when 24 h sampling is not feasible, the authors recommended that the first-void urine, representing accumulated overnight excretion, may provide the most consistent and representative alternative.^40^

If the urinary nucleic acid adductome is to be used as a biomarker of long-term, or indeed acute, exogenous or endogenous exposure, it is advantageous if the intra-individual (i.e., day-to-day or diurnal) variability is low compared to the inter-individual (i.e., person-to-person) variability. Under these conditions, a single sample can provide a reliable estimate of an individual’s typical exposure in epidemiologic studies. In contrast, high intra-individual variability introduces measurement error, reducing the accuracy of exposure classification and attenuating observed associations between exposure and health outcomes, as demonstrated in occupational exposure studies.^42^

Additionally, sex-specific differences may further contribute to possible diurnal variability in urinary nucleic acid adduct concentrations. Both males and females display distinct circadian patterns in hormone secretion, renal transporter activity, and xenobiotic metabolism, all of which influence urinary output and metabolite composition.^43–49^ Age also exerts a significant influence on diurnal urinary patterns in both sexes. Metabolomic studies have shown that urinary metabolites linked to mitochondrial energy metabolism and muscle mass decline differ markedly with age in both males and females.^50^ Functionally, maximum urine flow varies across age groups; younger men exhibit reduced peak flow rates during morning and nighttime hours compared to older individuals.^51^ Consequently, sex and age are also potential biological factors to consider in studies of the urinary adductome, as both the size and phase of adduct excretion may differ between males and females even under identical environmental conditions.

The objectives of the present study were to (1) assess whether there is any diurnal variation in the urinary excretion of nucleic acid adducts, (2) clarify the optimal technique for normalizing nucleic acid adduct concentrations in spot samples for urinary flow rate, and (3) assess the intra- and inter-individual variability in urinary nucleic acid adducts. Because both xenobiotic metabolism and kidney physiology exhibit known sex-dependent differences,^52–55^ analyses were performed separately for males and females, as well as in the combined population.

## Methods

### Urine samples

Urine samples were obtained from the Gothenburg Variability Biobank from Sweden. It includes a subsample of 60 healthy non-smokers, recruited in 2012 and 2013, mainly from employees or students at the Sahlgrenska Hospital and the University of Gothenburg.^39,56^ Repeated blood and urine samples from the full Swedish “variability biobank” are kept in a low temperature freezer and open for researchers examining normal variability of their favorite biomarkers. They provided timed urine samples at fixed times during two 24-h periods, approximately one week apart. The present study was performed using samples, stored at -80 °C, from 40 of these individuals, 20 men and 20 women, aged 21 – 64 y (mean 32 y). Mean body mass index was 23. Three timed urine samples from each individual were analyzed: a first void sample (obtained between 05.00 to 09.00 am) and an afternoon sample (obtained at approximately 03.00 pm) from day “A” to assess diurnal variability, and a first void sample from day “B”, one week later to evaluate day-to-day variability.

The urinary creatinine concentration was determined for each fresh urine sample using the Roche Creatinine enzymatic assay on the Cobas 6000 instrument (Roche Diagnostics Scandinavia AB, Solna, Sweden) with a limit of detection of 0.01 g/L, and traceable to the International Standardization with isotope dilution mass-spectrometry. Imprecision (coefficient of variation), as calculated by duplicate analyses, was 1.4– 3.6%. SG was measured with a Ceti Digit 012 refractometer (Medline, Oxfordshire, UK).

### Ethical approval

The study was approved by the Ethics Review Board at the University of Gothenburg (No. 213-12), and all participants signed a written informed consent.

### Urinary nucleic acid adductomics

A 0.5 mL aliquot of each urine sample was prepared for analysis using our dual phase SPE method,^57^ using an Isolute-ENV column followed by Hypersep-PHE. Samples were reconstituted with 100 µL of 1 mM aqueous ammonium acetate Sigma Aldrich (St. Louis, Massachusetts, USA). Each sample was spiked with a cocktail of isotopically labeled standards. This cocktail consisted of modified nucleobases (nBs), 2’-dNs, and ribonucleosides (rNs; Table S.1), sourced from LGC standards (North York, Canada).

Untargeted analysis was performed on a Thermo Scientific Vanquish Horizon HPLC system paired with an Orbitrap IQ X mass spectrometer. The acquisition method (Table S.2) was developed for the analysis of urinary adducts.^57^ This high-resolution MS/MS method captures a wide variety of adducts with no discrimination against any type of adduct. The HPLC gradient comprised H_2_O, methanol, and formic acid, and ammonium acetate all obtained from Fisher Chemical (Fair Lawn, New Jersey, USA).

The post-acquisition analysis was performed using a combination of MSconvert, FeatureHunter, MZ mine and Metaboanalyst. Raw data were converted to ‘.mzml’ format with MSconvert. Once converted, untargeted adductomics analysis was achieved using FeatureHunter^58^ , in which a series of fragmentation patterns characteristic to nucleic acid adducts was used to identify MS/MS signals that could be from adducts. In addition, the ‘mzml’ converted data were processed through MZmine, which filters the raw data and removes any noise. The MZmine and FeatureHunter outputs were then merged, and the overlapping signals were regarded as adducts. With FeatureHunter removing unnecessary data from MZmine and, conversely, MZmine removing noise from FeatureHunter the total urinary adductome was revealed.

More specifically, FeatureHunter allows for untargeted analysis based on characteristic fragmentation patterns derived from the chemical structure of nucleic acids. Each chemical class of nucleic acid adduct was searched for based on the following fragmentation patterns. As previously stated, 2’-dNs are characterized by the neutral loss (NL) of 116.0473 *m/z* associated with a sugar loss. The nBs are more complex as there are multiple types of nBs, each thymine, cytosine, guanine, adenine, and uracil have their own associated NL, respectively: 126.0429 *m/z*, 111.043 *m/z*, 151.0494 *m/z*, 135.0544 *m/z*, and 112.0272 *m/z* for each nBs. Finally, rNs are characterized by the NL of 132.0422 *m/z* associated with the loss of a ribose sugar, additionally a NL of 146.0579 *m/z* is associated with methylated ribonucleosides. Any signal containing either NL were considered to be a rN.

The nucleic acid adducts discovered through this untargeted approach were then normalized for either creatinine or specific gravity (see above). For each urine sample, creatinine normalization was achieved by taking the signal intensity (determined by the Orbitrap IQX) or concentration (as calculated comparing sample signal intensity to isotopically labeled standard signal intensity) of the adducts and dividing this by the creatinine concentration (g/L).^59,60^ For each urine sample, to normalize for specific gravity, the study populations’ average specific gravity (SG is unitless) was compared to the individual’s specific gravity to generate an adjustment factor which was then used to adjust the signal intensity or concentration of the adducts.^61^ After the levels of adducts were normalized for either creatinine or specific gravity, statistical analyses were performed using Metaboanalyst.

In addition to untargeted analysis, three adducts were selected to undergo targeted analysis which allowed us to perform the inter- and intra-individual variability studies. One adduct from each chemical species 8-oxodG (2’dN), 7-Me-Gua (nB), and 2-Me-Cyt (rN) was selected from the list of standards (Table S.1) for targeted analysis. For this targeted analysis the concentration of each target was determined using a one-point internal calibration to the known concentration of the equivalent isotopically labeled standard at 5 ppm. Once the concentration at each time point was established, the excretion rate was determined by comparing the mass of adduct (concentration x volume) found in the total void to the time between voids.

### Statistical analysis

Variability in urinary excretion was determined by two methods, the first of which was calculating the coefficient of variation (CV). The CV describes only the intra-individual variability as the ratio between the SD and the mean of replicate samples. The other method was to simultaneously calculate the intra- and inter-individual variance and the intraclass correlation (ICC). The ICC is the ratio of the inter-individual variance to the total (sum of inter- and intra-individual) variance calculated on ln-transformed data. A high ICC is preferable since then the variability between individuals is larger than the variability within individuals over time. Partitioning of intra- and inter-individual variability was performed using the procedure Nested in SAS version 9.4 (SAS Institute, Cary, NC, USA).

Within the averaged adductome maps, lower average signal intensities generally reflect features detected in a subset of individuals, whereas higher-intensity signals indicate adducts more consistently observed across the cohort. For downstream statistical analyses, adductomes from individual participants were grouped according to relevant biological variables, including time of collection, sex, and age. To improve robustness and reduce the influence of sparsely detected features, only adducts present in at least 50% of individuals were retained for analysis using MetaboAnalyst. Subsequent differential analyses, including volcano plots, therefore represent a filtered subset of the total detected adductome, focusing on features consistently observed across the population to enable reliable comparison between first void and afternoon samples.

## Results

Individual urine samples were analyzed using an untargeted nucleic acid adductomics workflow to define subject-specific urinary nucleic acid adductomes. Figure 1A illustrates the nucleic acid adducts present in a representative individual’s 03.00 pm spot urine sample. The adduct map shows the individual’s adduct type distribution and relative abundance. To facilitate chemical classification, the adducts were stratified into three primary categories: nucleobases (119 nB; Figure 1B), ribonucleosides (44 rN; Figure 1C), 2’-deoxyribonucleosides (9 2’-dN; Figure 1D), of which the number of nB adducts predominated.^30,62–64^ A selection of the signals recovered is from the isotopically labeled cocktail. These were marked with identity and asterisk to indicate their isotopic label. It should be noted that across the grouping of chemical structures there is an overlapping of adduct signals. This is due to overlapping fragmentation patterns that may indicate the identity of more than one chemical structure.

**Figure 1.**
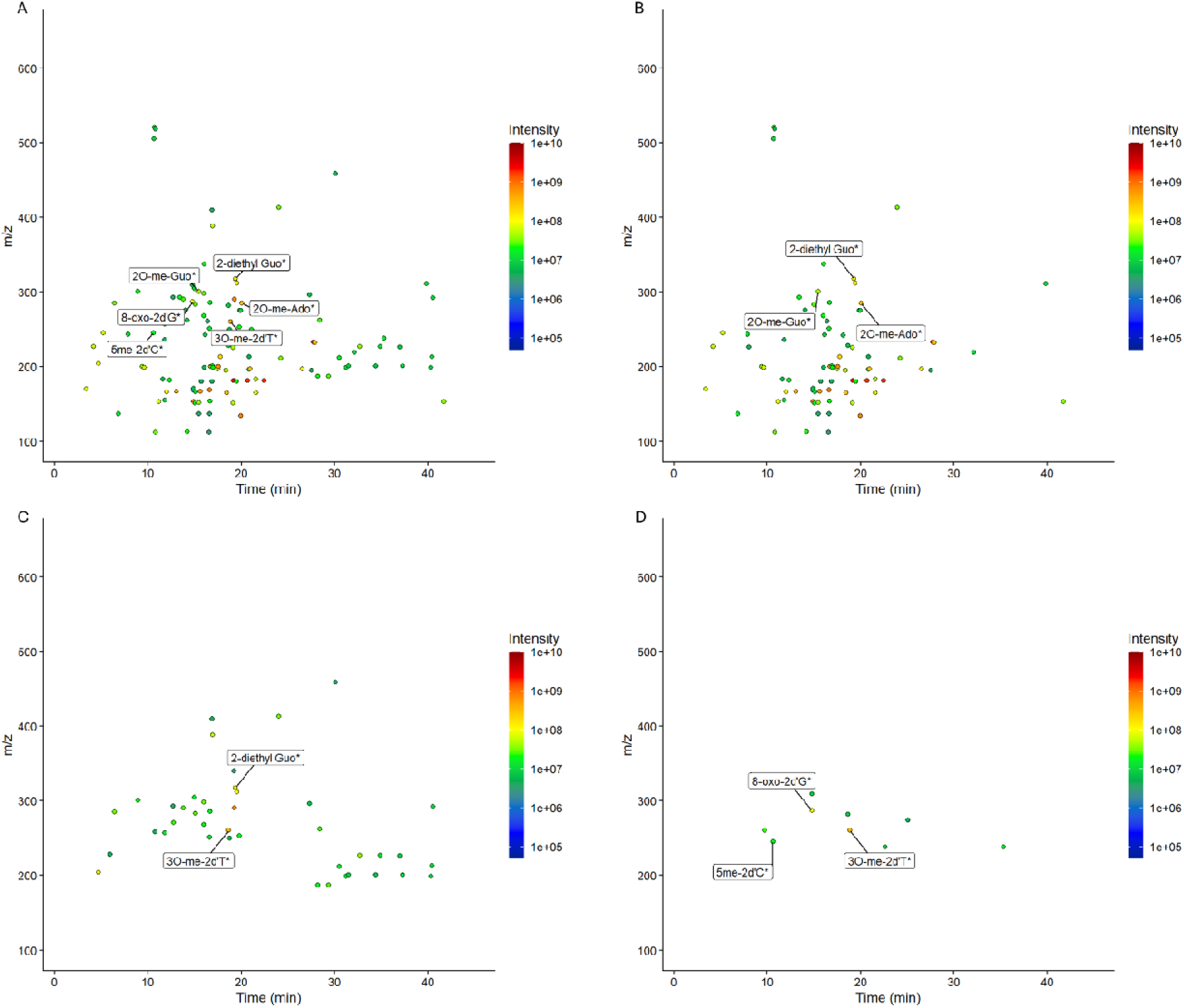
(A) A representative urinary nucleic acid adductome map illustrating the breadth and intensity of nucleic acid adducts. The urinary nucleic acid adductome was then stratified into (B) nB, (C) rN, and (D) 2’-dN adducts. These urinary nucleic acid adductome maps were derived from a spot urine sample from one individual taken at the 03.00 pm sampling time, totaling 130 signals across 2’-dN, nB and rN. nB adducts predominate in the urine, compared to both rN and 2’-dN.

### Intra- and inter-individual variability in the urinary concentration of representative nucleic acid adducts

Targeted analysis determined that the unadjusted mean concentrations of the representative adducts, 8-oxodG (2’-dN), 7-Me-Gua (nB), and 2-Me-Cyt (rN) were 1.35 nM, 9.34 nM, and 8.17 nM, respectively (Table 3), are consistent with previous reports.^65–68^ In comparing first void samples to afternoon void, distinct temporal patterns emerged which remained even following normalization to creatinine or SG. For 8-oxodG, the highest concentration was observed in the afternoon sample (1.63 nM), and this remained the case regardless of the normalization method. In contrast, unadjusted concentrations of both 7-Me-Gua and 2-Me-Cyt were highest in the second first-void collection (10.65 nM and 9.70 nM, respectively); however, after adjustment for diuresis, using creatinine, peak concentrations were then seen in the afternoon void. It should be noted that none of these adducts were present at statistically significantly higher levels in either the first or afternoon void.

The results for excretion rate further supported these observations where all three adducts, excretion rates were elevated in afternoon samples relative to first-void collections. Specifically, 8-oxodG exhibited an average excretion rate of 0.17 nM/h (afternoon) compared to 0.081 and 0.049 nM/h in the two first-void samples (one sample from each of the two weeks). Similarly, 7-Me-Gua showed mean excretion rates of 0.68 nM/h (afternoon) versus 0.45 and 0.42 nM/h (first void), and 2-Me-Cyt showed rates of 0.53 nM/h versus 0.37 and 0.40 nM/h for afternoon versus first-void samples, respectively (Table 1). Notably, excretion rates between first-void samples collected one week apart were comparable across the three adducts analyzed, indicating temporal stability in baseline adduct elimination despite differences between the collection dates. Intra-individual variability in repeated first-void samples showed CVs for unadjusted concentrations of 0.33 for 8-oxo-dG, 0.46 for 7-Me-Gua, and 0.42 for 2-Me-Cyt (Table 1). The impact of adjustment for creatinine or SG was not evident, although the CV increased for 8-oxodG but decreased for 7-Me-Gua and 2-Me-Cyt (Table 1). The inter-individual variability was higher than the intra-individual variability only for creatinine- and SG-adjusted 8-oxodG. For unadjusted concentrations, ICC values were below 0.40 for all adducts, indicating limited reliability of single measurements for epidemiologic applications.^69^ For 8-oxodG, ICC values were 0.58 (creatinine) and 0.55 (SG). ICC values for 7-Me-Gua and 2-Me-Cyt were low regardless of adjustments suggesting that for these urinary adducts quantitative applications may require repeated sampling. It must be noted that this is an analysis of variation in a targeted application.

**Table 1.** A single spot sample of urinary 8-oxodG concentration, adjusted for creatinine or SG, is useful as a biomarker when comparing populations. The excretion rates of three adducts 8-oxodG, 7-Me-Gua and 2-Me-Cyt (representative of the three classes of urinary nucleic acid adduct: 2’-dN, nB and rN) are higher in the afternoon sample compared to the morning sample. For all individuals (n = 40) the excretion rate was calculated by taking the measured concentration of adduct in urine and dividing it by the time between voids. Intra- and inter-individual variability of the log-transformed data was calculated for the two first void samples which were provided one week apart.

|  | **Correction Factor** | **8-oxodG** | **7-Me-Gua** | **2-Me-Cyt** |
| --- | --- | --- | --- | --- |
| **First void (Day 1) concentration (nM)** | Unadjusted | 1.42 | 8.46 | 7.47 |
|  | Creatinine-adjusted | 1.61 | 8.25 | 6.76 |
|  | SG-adjusted | 1.65 | 8.65 | 7.15 |
| **First void (Day 7) concentration (nM)** | Unadjusted | 1.02 | 10.65 | 9.70 |
|  | Creatinine-adjusted | 1.07 | 9.01 | 7.41 |
|  | SG-adjusted | 1.14 | 10.21 | 8.54 |
| **Afternoon sample (Day 1) concentration (nM)** | Unadjusted | 1.63 | 8.92 | 7.34 |
|  | Creatinine-adjusted | 2.97 | 10.41 | 7.79 |
|  | SG-adjusted | 2.63 | 10.15 | 7.76 |
| **Excretion Rate (nM/h)** | First Void (A) | 0.081 | 0.42 | 0.37 |
|  | First Void (B) | 0.049 | 0.45 | 0.40 |
|  | Afternoon (A) | 0.17 | 0.68 | 0.53 |
| **Intra-individual variability CV** | Un corrected | 0.33 | 0.46 | 0.42 |
|  | Creatinine-adjusted | 0.53 | 0.34 | 0.24 |
|  | SG-adjusted | 0.43 | 0.39 | 0.27 |
| **Intra-Individual variability ratio to total variability** | Un corrected | 0.62 | 0.71 | 0.63 |
|  | Creatinine-adjusted | 0.42 | 0.85 | 0.80 |
|  | SG-adjusted | 0.45 | 0.78 | 0.86 |
| **Inter- individual variability ratio to total variability = intra-class correlation (ICC)** | Un corrected | 0.38 | 0.29 | 0.37 |
|  | Creatinine-adjusted | 0.58 | 0.15 | 0.20 |
|  | SG-adjusted | 0.55 | 0.22 | 0.14 |

### Variability in the untargeted urinary nucleic acid adductome

Adductome maps are composed of two components, (i) the range of adducts, and (ii) the intensity of these adducts. We examined diurnal variation in each adduct by comparing the variation in intensity of each adduct across sampling times represented as a volcano plots, using paired analysis to ensure individual variability was taken into consideration. Regardless of adjustment method (i.e., unadjusted, Figure 2A; creatinine-adjusted, Figure 2B; and SG-adjusted, Figure 2C) the ratios between adduct intensities in first void and afternoon samples were present in significantly higher intensities in the first void samples (logFC < -1 in Figure 2) with only two adducts showing a statistically significant increase in the afternoon void.

**Figure 2.**
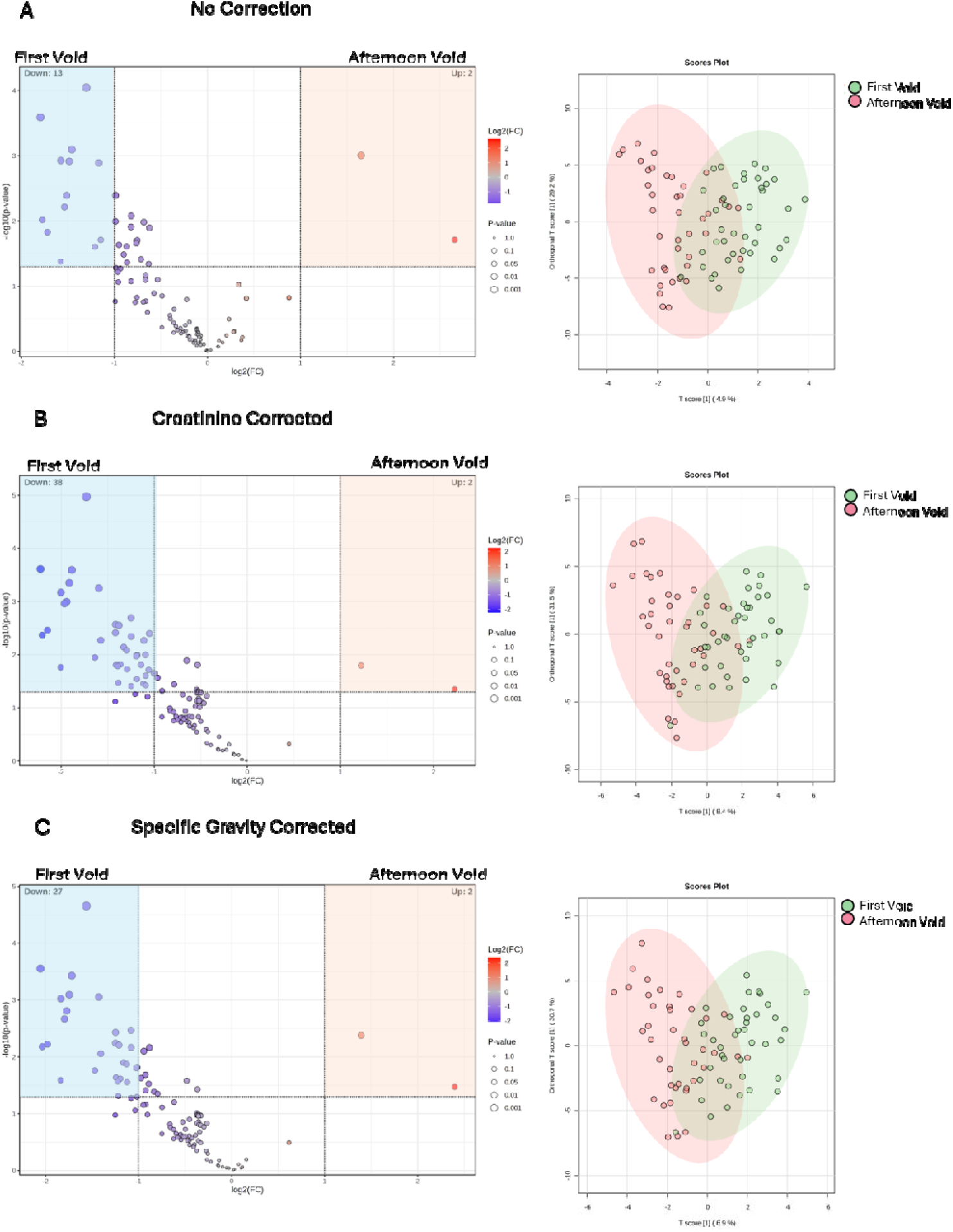
There was minimal variation in urinary nucleic acid adductome between first void and afternoon samples when (A) unadjusted, (B) adjusted for creatinine, or (C) SG. Those adducts that are represented in blue to the left of x-axis -1 and above y axis 1 decreased significantly from the first void to the afternoon void. The majority of the adducts were significantly higher in intensity in the first void samples, with only two adducts being significantly higher in the afternoon void. In addition, multivariate OPLS-DA was performed in MetaboAnalyst to compare the total adductome profiles between first void and afternoon void in individual urine samples (n=40). The trend remained, as the score plots showed very limited separation between time points. Note that the OPLS score plots have no “direction” but only shows the separation between time points.

Additionally, each individual sample underwent untargeted analysis to determine difference in the total adductome. Multivariate analysis using orthogonal partial least squares–discriminant analysis (OPLS-DA) was performed to evaluate global differences in urinary adductomes between first void and afternoon samples (Figure 2). OPLS-DA is a supervised analysis strategy that will separate predefined groups by filtering out nonsignificant features to highlight the differences driven by significant features. Across all normalization strategies (unadjusted, creatinine-adjusted, and SG–adjusted), multivariate score plots demonstrated minimal cluster separation between sampling time points, indicating negligible change in the composition of the urinary adductome over a single day. The majority of the clusters overlapped indicating minimal differences between sampling time, supporting the results from the volcano analysis, while the primary predictive component (time of sampling, T score [1]), the variable being tested for, was low and comparable across models (4.9 %, 8.4 %, and 6.9 % for unadjusted, creatinine-, and SG-corrected data, respectively). In contrast, a greater proportion of cluster spread was captured by the orthogonal components (29.2 %, 31.5 %, and 30.7 %, respectively), variables unaccounted for, suggesting that most variability reflects intra-group heterogeneity not associated with time of sampling. Consistent with this interpretation, score plots exhibited elongated, vertically distributed clusters with substantial overlap between time points, indicating poor discrimination between time points regardless of normalization approach. Targeted investigation of adducts listed in Table 1 (8-oxodG, 7-Me-Gua, 2-Me-Cyt) revealed that although these adducts were recovered at a higher concentration in afternoon sampling, these differences were not statistically significant.

All subsequent analyses were performed using SG–corrected data, due to the previously reported variability of creatinine.^70^ Across the cohort of 40 participants, SG-adjusted data revealed 27 adducts (Table 1) that were significantly different in their relative signal intensity between first-void and afternoon samples [a combination of p < 0.05 and log(Fold change <-1 or >1)] (Figure 3). At the population level, signal intensities of 2’-dN, nB, and rN adducts were generally elevated in the samples collected at first void, with those that were statistically higher in their signal al present in the first void sample, adduct signal intensities did change relative to diurnal variation with higher concentrations present in the first void sample (Figure 3A). However, when stratified by sex, the magnitude of change differed between groups. Males exhibited a smaller number of adducts (eight adducts; Figure 3B) that significantly varied by time than females (39 significantly different adducts across time points; Figure 3C). All present at significantly higher levels in morning samples than afternoon void samples. Multivariate OPLS-DA analysis was consistent with the observed patterns from the volcano plots. These findings indicate that inter-individual variability (i.e., metabolic turnover, age, lifestyle, etc.), rather than diurnal variation, was the predominant source of variation within this dataset, regardless of cluster separation since the predictive variance was low compared to the orthogonal component.

**Figure 3.**
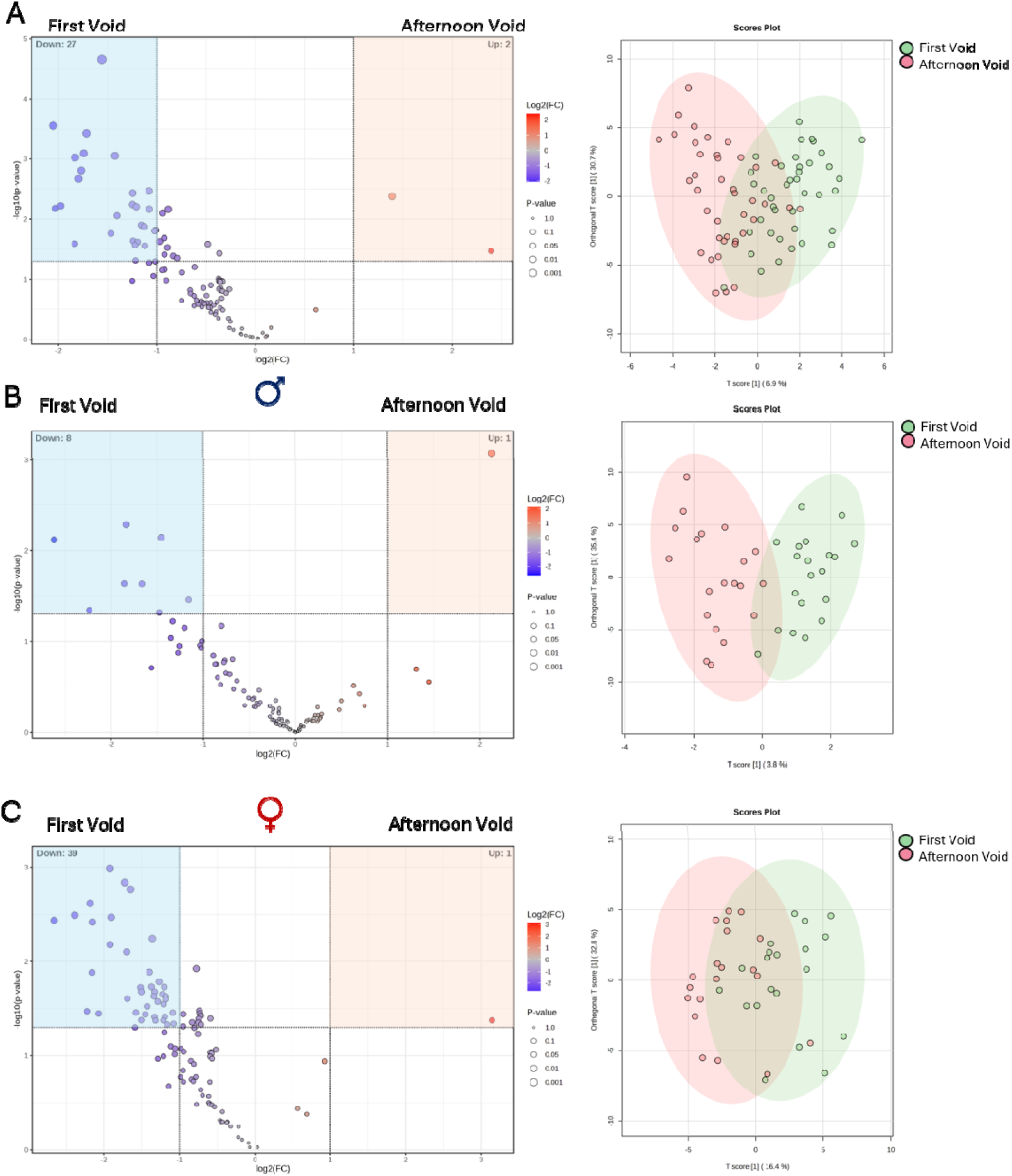
The adducts present with significantly higher intensities were mainly present in the first void sample with only one or two adducts higher in the afternoon sample. In each (A) the whole population, (B) males, and (C) females. Those adducts that are represented in blue to the left of x-axis -1 and above y axis 1 decreased significantly from the first void to the afternoon void. In the whole population only 27 adduct intensities were significantly higher in the first void sample, in males this was true for eight adducts, and for 39 of the adducts in females. When multivariate OPLS-DA was performed in MetaboAnalyst to compare the total adductome profiles between first void and afternoon void in individual urine samples score plots showing very limited separation between time points, the trend remained for (A) the whole population (B) males, and (C) females.

To determine whether the total number of adducts (i.e. the range of adducts), inclusive of all class types, changed by time point of sampling or sex, adduct counts were compared across these factors. At the population level, the total number of adducts excreted did not fluctuate with diurnal cycling (Figure 4). This analysis reflects the number of adducts detected, not accounting for differences in signal intensity. Stratification by sex yielded similar results. In females, urine samples collected at first void, 3 pm, and first void one week later contained an average of 153, 153, and 170 adducts, respectively, with no significant differences across time points. Similarly, in males, mean adduct counts were 144, 169, and 197 at the corresponding intervals, with no significant variation in the number of adducts present.

**Figure 4.**
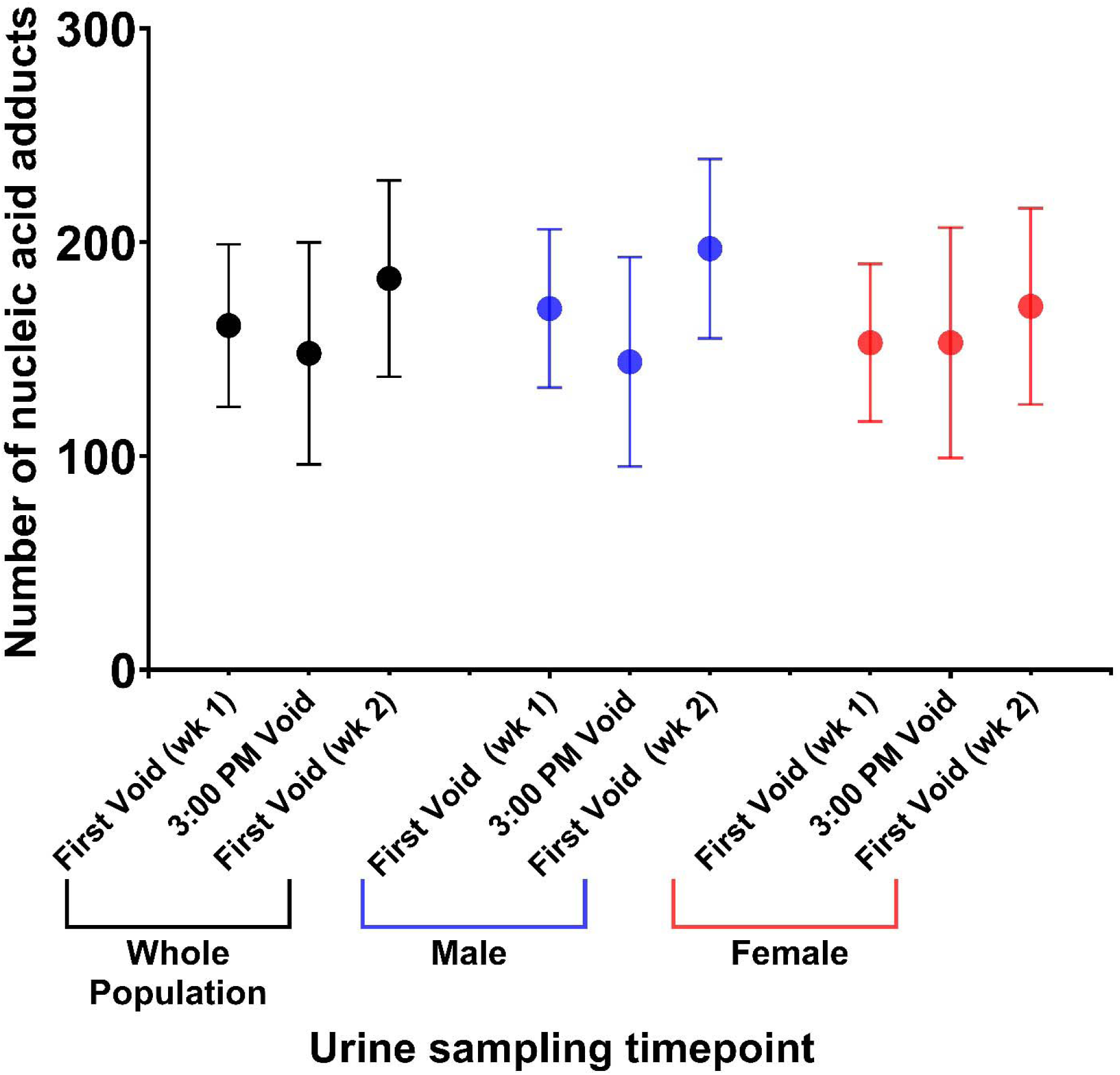
The number of adducts comprising the urinary nucleic acid adductome were independent of diurnal variation. There was no statistically significant difference in the number of adducts measured in the 03.00 pm sample compared to the two samples collected at first void (separated by one week). This trend was also present when the population was segregated by sex. Note that this is not a quantitative measure of the adduct intensity and only reflects the number of adducts recovered in an individual’s samples. This demonstrates that qualitative analysis may be accomplished with a spot urine sample obtained at any timepoint.

### Effect of sex and age on urinary nucleic acid adductome

We also examined the possible effect of sex and age, irrespective of the time of sampling. When SG-adjusted first-void samples from males and females (n = 20 per group) were compared, there was no overall difference in signal intensity, with the fold change centered around 1. Only four adducts were present in statistically significantly higher amounts in males (p < 0.05) (Figure 5A). Due to the limited group size, we were unable to exclude subtle sex-dependent differences in adduct excretion, metabolic turnover or differences in the exposures of each sex.^53^ OPLS-DA plots depict slightly merged elongated clusters with a primary predictive component explaining 9.4% and the orthogonal component explaining 37.3% of differences. These results indicate that sex differences are statistically apparent but contribute minimally towards separation, where other variables may have greater contributions. For this reason, age was selected as another key variable that can drive metabolic differences in adduct excretion.

**Figure 5.**
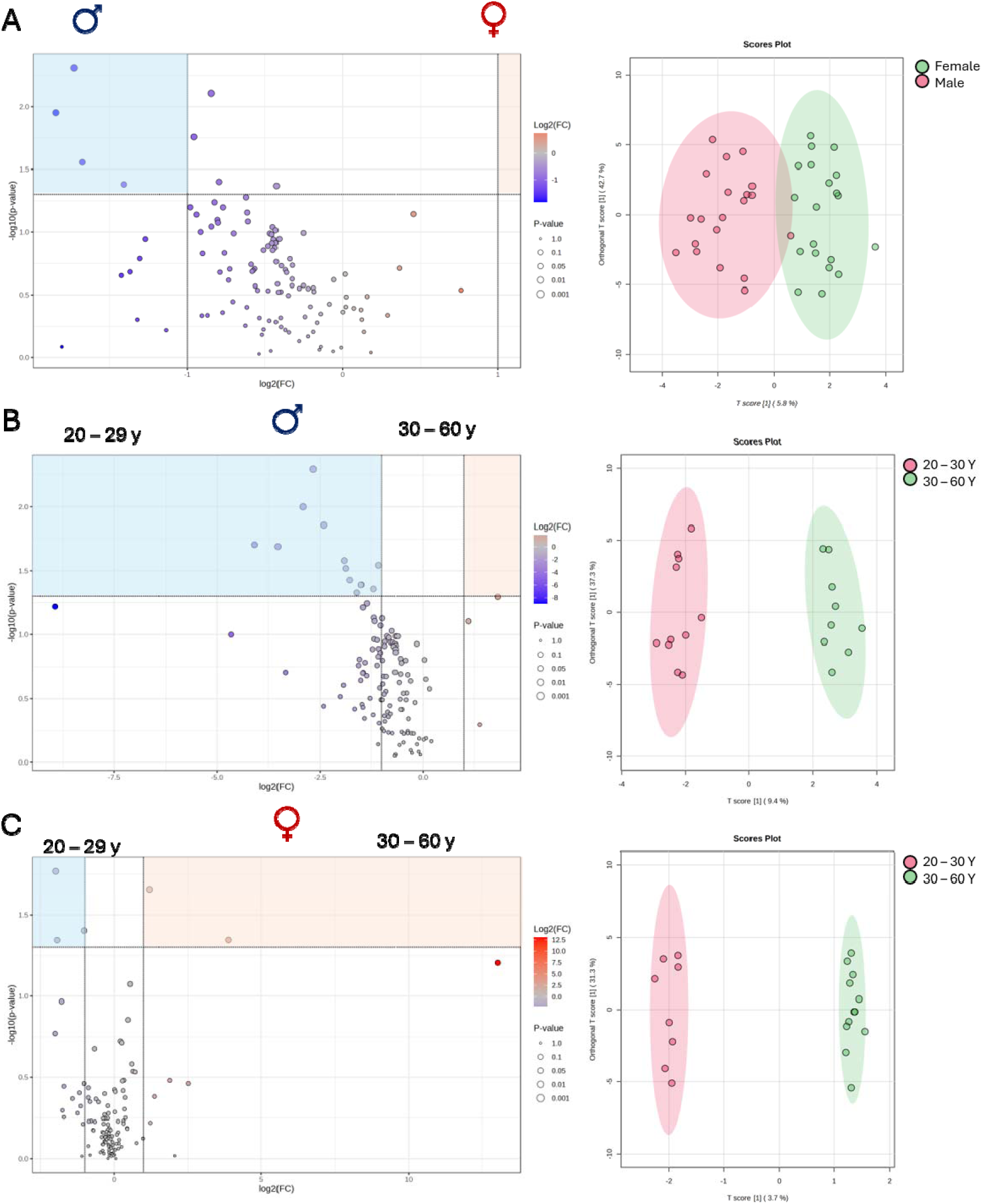
Sex and age are contributing factors to the variation of the urinary nucleic acid adductome. (**A**) Adduct intensities are higher in urine samples from males compared to females. When stratifying by both sex and age, in males **(B)** adduct intensities were higher in the 21 – 29 y group. In females **(C)** two adducts had higher intensities in the 21 – 29 y group and two adducts in the 30 – 60 y age group. The multivariate OPLS-DA showed some separation between (A) males and females and (B) between age groups in males. In females, the separation by age groups (x-axis) was small (4%) compared to the orthogonal component (31%).

Age-related differences in urinary adduct excretion profiles were evaluated by dividing participants into two age groups: 21–29 years and 30–60 years. In males, all adducts were present at significantly higher intensity in the younger population (21-29 years), compared to the older population (Figure 5 B). In contrast, in females there were two adducts (Table 2) that were of significantly higher intensity in the younger population and two that were significantly higher in the older population (Figure 5 C).

**Table 2.**
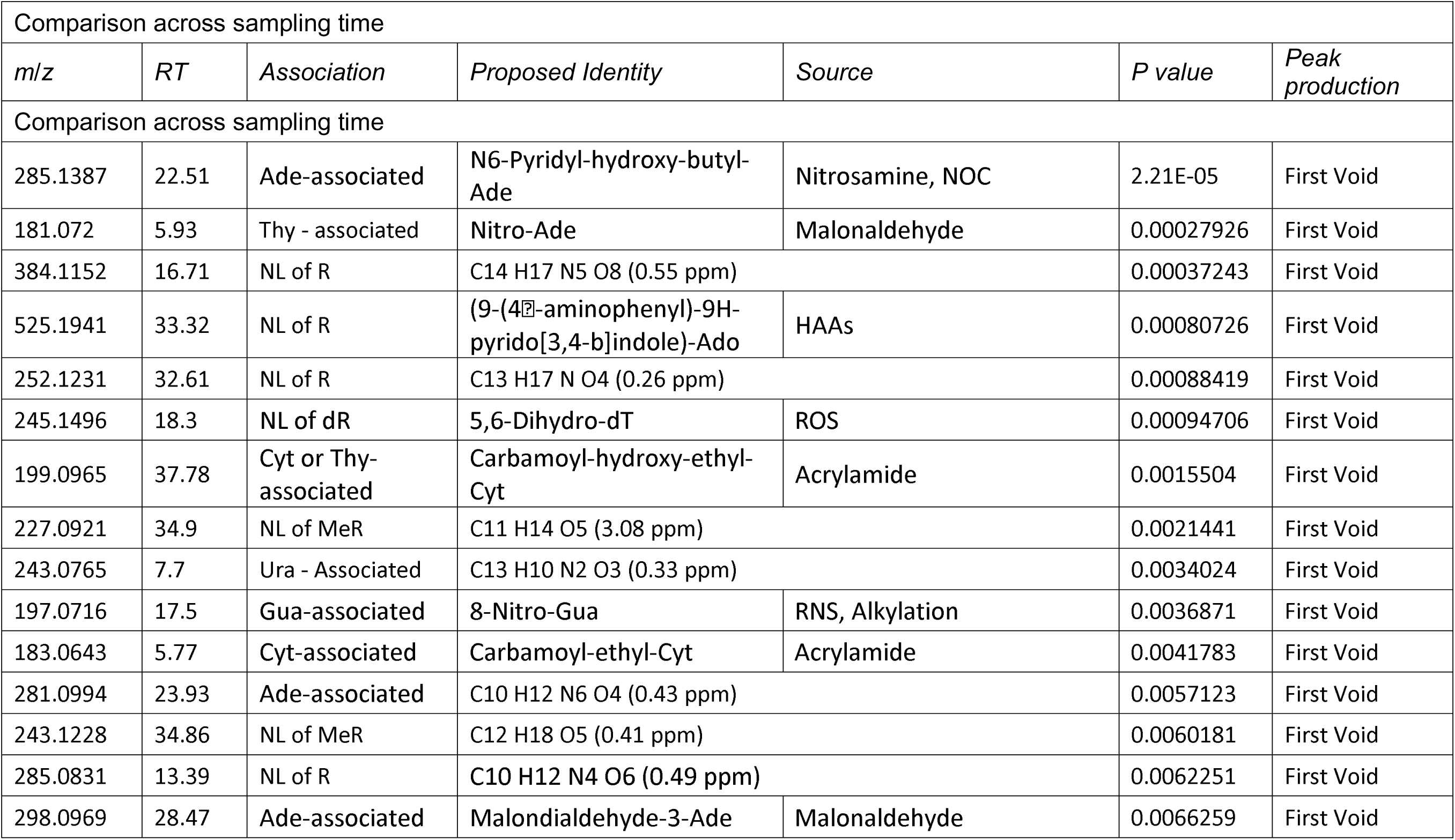
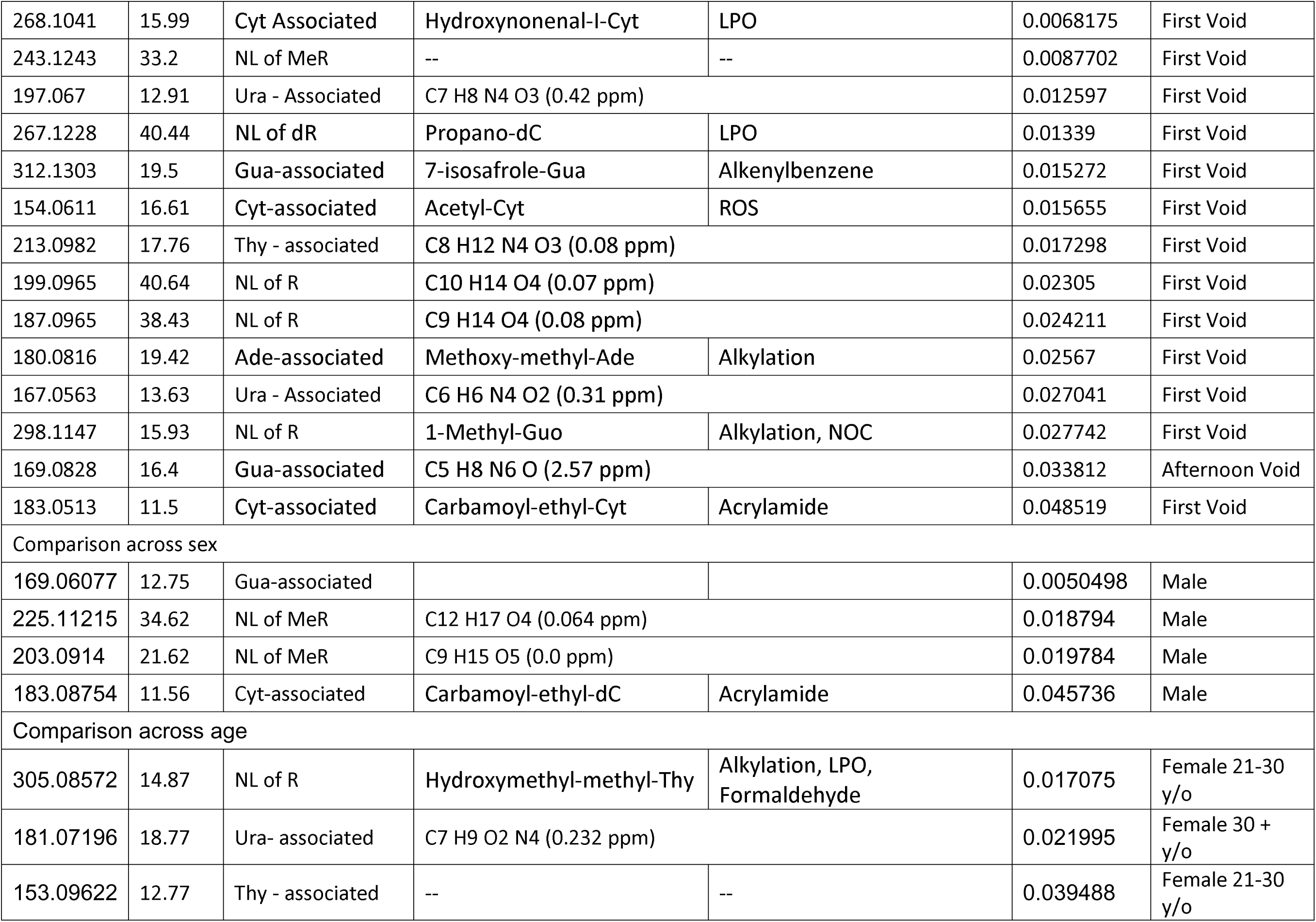
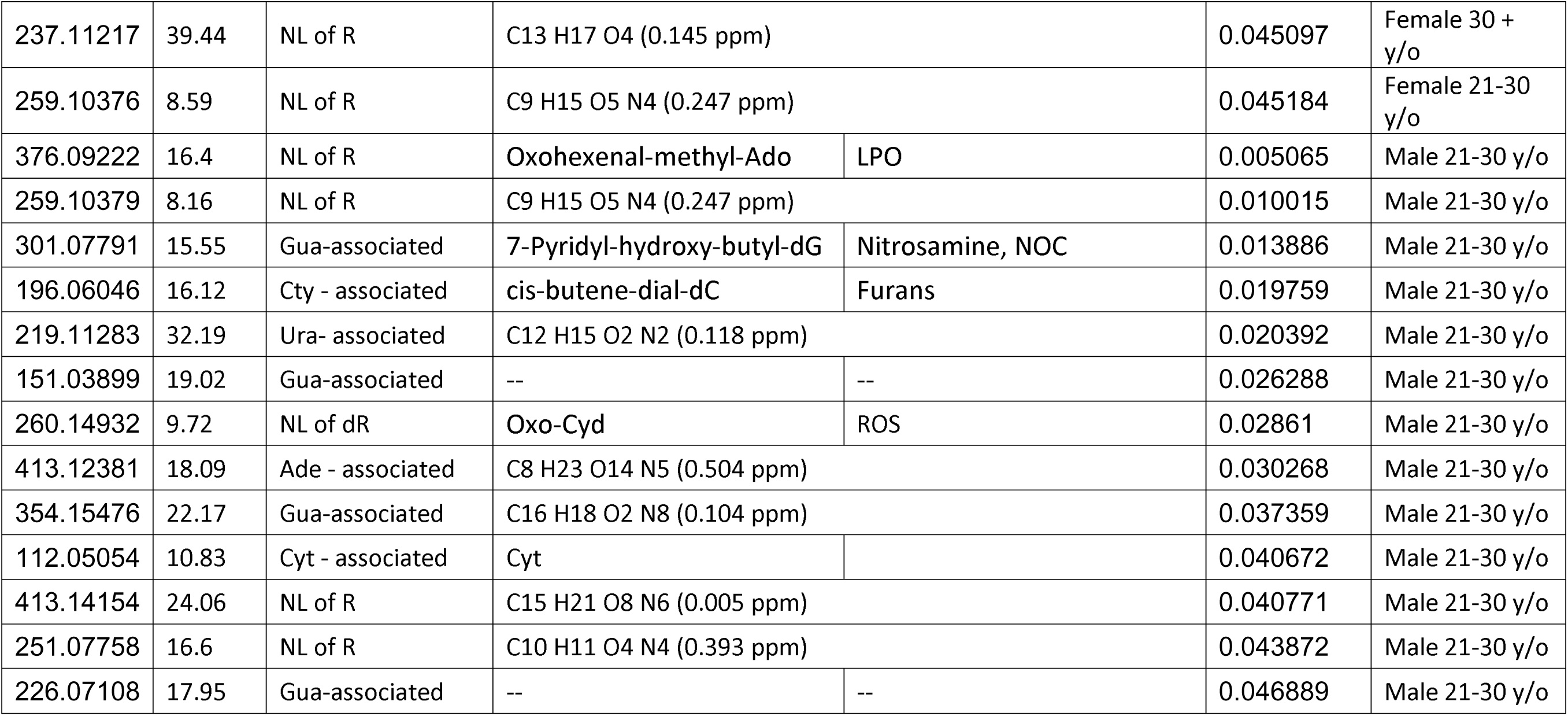
Putative identities of the adducts whose intensities significantly changed across sampling times, sex and age. Based on their characteristic fragmentation patterns, the following chemical formulae were proposed. A selection of these adducts were identified using the Stanstrup et al., database.^91^ Each proposed chemical formula is within 5 ppm of the measured m/z ensuring the proposed formula was congruent with the accurate mass measured. Those with no formula were unidentifiable based on the characteristic fragments and the 5 ppm requirement for the chemical formula.

| Comparison across sampling time |  |  |  |  |  |  |
| --- | --- | --- | --- | --- | --- | --- |
| *m/z* | *RT* | *Association* | *Proposed Identity* | *Source* | *P value* | *Peak production* |
| Comparison across sampling time |  |  |  |  |  |  |
| 285.1387 | 22.51 | Ade-associated | N6-Pyridyl-hydroxy-butyl-Ade | Nitrosamine, NOC | 2.21E-05 | First Void |
| 181.072 | 5.93 | Thy - associated | Nitro-Ade | Malonaldehyde | 0.00027926 | First Void |
| 384.1152 | 16.71 | NL of R | C14 H17 N5 O8 (0.55 ppm) |  | 0.00037243 | First Void |
| 525.1941 | 33.32 | NL of R | (9-(4-aminophenyl)-9H-pyrido[3,4-b]indole)-Ado | HAAs | 0.00080726 | First Void |
| 252.1231 | 32.61 | NL of R | C13 H17 N O4 (0.26 ppm) |  | 0.00088419 | First Void |
| 245.1496 | 18.3 | NL of dR | 5,6-Dihydro-dT | ROS | 0.00094706 | First Void |
| 199.0965 | 37.78 | Cyt or Thy-associated | Carbamoyl-hydroxy-ethyl-Cyt | Acrylamide | 0.0015504 | First Void |
| 227.0921 | 34.9 | NL of MeR | C11 H14 O5 (3.08 ppm) |  | 0.0021441 | First Void |
| 243.0765 | 7.7 | Ura - Associated | C13 H10 N2 O3 (0.33 ppm) |  | 0.0034024 | First Void |
| 197.0716 | 17.5 | Gua-associated | 8-Nitro-Gua | RNS, Alkylation | 0.0036871 | First Void |
| 183.0643 | 5.77 | Cyt-associated | Carbamoyl-ethyl-Cyt | Acrylamide | 0.0041783 | First Void |
| 281.0994 | 23.93 | Ade-associated | C10 H12 N6 O4 (0.43 ppm) |  | 0.0057123 | First Void |
| 243.1228 | 34.86 | NL of MeR | C12 H18 O5 (0.41 ppm) |  | 0.0060181 | First Void |
| 285.0831 | 13.39 | NL of R | C10 H12 N4 O6 (0.49 ppm) |  | 0.0062251 | First Void |
| 298.0969 | 28.47 | Ade-associated | Malondialdehyde-3-Ade | Malonaldehyde | 0.0066259 | First Void |
| 268.1041 | 15.99 | Cyt Associated | Hydroxynonenal-l-Cyt | LPO | 0.0068175 | First Void |
| 243.1243 | 33.2 | NL of MeR | -- | -- | 0.0087702 | First Void |
| 197.067 | 12.91 | Ura - Associated | C7 H8 N4 O3 (0.42 ppm) |  | 0.012597 | First Void |
| 267.1228 | 40.44 | NL of dR | Propano-dC | LPO | 0.01339 | First Void |
| 312.1303 | 19.5 | Gua-associated | 7-isosafrole-Gua | Alkenylbenzene | 0.015272 | First Void |
| 154.0611 | 16.61 | Cyt-associated | Acetyl-Cyt | ROS | 0.015655 | First Void |
| 213.0982 | 17.76 | Thy - associated | C8 H12 N4 O3 (0.08 ppm) |  | 0.017298 | First Void |
| 199.0965 | 40.64 | NL of R | C10 H14 O4 (0.07 ppm) |  | 0.02305 | First Void |
| 187.0965 | 38.43 | NL of R | C9 H14 O4 (0.08 ppm) |  | 0.024211 | First Void |
| 180.0816 | 19.42 | Ade-associated | Methoxy-methyl-Ade | Alkylation | 0.02567 | First Void |
| 167.0563 | 13.63 | Ura - Associated | C6 H6 N4 O2 (0.31 ppm) |  | 0.027041 | First Void |
| 298.1147 | 15.93 | NL of R | 1-Methyl-Guo | Alkylation, NOC | 0.027742 | First Void |
| 169.0828 | 16.4 | Gua-associated | C5 H8 N6 O (2.57 ppm) |  | 0.033812 | Afternoon Void |
| 183.0513 | 11.5 | Cyt-associated | Carbamoyl-ethyl-Cyt | Acrylamide | 0.048519 | First Void |
| Comparison across sex |  |  |  |  |  |  |
| 169.06077 | 12.75 | Gua-associated |  |  | 0.0050498 | Male |
| 225.11215 | 34.62 | NL of MeR | C12 H17 O4 (0.064 ppm) |  | 0.018794 | Male |
| 203.0914 | 21.62 | NL of MeR | C9 H15 O5 (0.0 ppm) |  | 0.019784 | Male |
| 183.08754 | 11.56 | Cyt-associated | Carbamoyl-ethyl-dC | Acrylamide | 0.045736 | Male |
| Comparison across age |  |  |  |  |  |  |
| 305.08572 | 14.87 | NL of R | Hydroxymethyl-methyl-Thy | Alkylation, LPO, Formaldehyde | 0.017075 | Female 21-30 y/o |
| 181.07196 | 18.77 | Ura- associated | C7 H9 O2 N4 (0.232 ppm) |  | 0.021995 | Female 30 + y/o |
| 153.09622 | 12.77 | Thy - associated | -- | -- | 0.039488 | Female 21-30 y/o |
| 237.11217 | 39.44 | NL of R | C13 H17 O4 (0.145 ppm) |  | 0.045097 | Female 30 + y/o |
| 259.10376 | 8.59 | NL of R | C9 H15 O5 N4 (0.247 ppm) |  | 0.045184 | Female 21-30 y/o |
| 376.09222 | 16.4 | NL of R | Oxohexenal-methyl-Ado | LPO | 0.005065 | Male 21-30 y/o |
| 259.10379 | 8.16 | NL of R | C9 H15 O5 N4 (0.247 ppm) |  | 0.010015 | Male 21-30 y/o |
| 301.07791 | 15.55 | Gua-associated | 7-Pyridyl-hydroxy-butyl-dG | Nitrosamine, NOC | 0.013886 | Male 21-30 y/o |
| 196.06046 | 16.12 | Cty - associated | cis-butene-dial-dC | Furans | 0.019759 | Male 21-30 y/o |
| 219.11283 | 32.19 | Ura- associated | C12 H15 O2 N2 (0.118 ppm) |  | 0.020392 | Male 21-30 y/o |
| 151.03899 | 19.02 | Gua-associated | -- | -- | 0.026288 | Male 21-30 y/o |
| 260.14932 | 9.72 | NL of dR | Oxo-Cyd | ROS | 0.02861 | Male 21-30 y/o |
| 413.12381 | 18.09 | Ade - associated | C8 H23 O14 N5 (0.504 ppm) |  | 0.030268 | Male 21-30 y/o |
| 354.15476 | 22.17 | Gua-associated | C16 H18 O2 N8 (0.104 ppm) |  | 0.037359 | Male 21-30 y/o |
| 112.05054 | 10.83 | Cyt - associated | Cyt |  | 0.040672 | Male 21-30 y/o |
| 413.14154 | 24.06 | NL of R | C15 H21 O8 N6 (0.005 ppm) |  | 0.040771 | Male 21-30 y/o |
| 251.07758 | 16.6 | NL of R | C10 H11 O4 N4 (0.393 ppm) |  | 0.043872 | Male 21-30 y/o |
| 226.07108 | 17.95 | Gua-associated | -- | -- | 0.046889 | Male 21-30 y/o |

Multivariate analysis using OPLS-DA was performed to assess differences in class-specific adduct intensities across the full cohort stratified by sex (n = 20 per group) and age (21–30 vs >30 years). Stratification by sex revealed minimal separation across all adduct classes. Variance was predominantly captured by orthogonal components (42.7%), with comparatively low contribution from the predictive components (5.8%), indicating that within-group variability exceeded sex-dependent differences.

### Putative identification of nucleic acid adducts exhibiting variation

To further explore the possibility of individual adducts as potential biomarkers, those showing significant temporal or group-dependent variation were cross-referenced against the Stanstrup^71^ database for putative identification. Of the 132 adducts that were found in the urinary samples from this population, there were a total of 51 adducts that significantly varied over the comparisons made here. Of those 51, 22 could be identified using a database previously reported in literature.^71^ The adducts identified at significantly higher levels in the first void sampling when comparing sampling times were N6-Pyridyl-hydroxy-butyl-Ade, Nitro-Ade, (9-(4′-aminophenyl)-9H-pyrido[3,4-b]indole)-Ado, 5,6-Dihydro-dT, Carbamoyl-hydroxy-ethyl-Cyt, 8-Nitro-Gua, Carbamoyl-ethyl-Cyt, Malondialdehyde-3-Ade, Hydroxynonenal-I-Cyt, Propano-dC, 7-isosafrole-Gua, Acetyl-Cyt, Methoxy-methyl-Ade, 1-Methyl-Guo, and Carbamoyl-ethyl-Cyt (Table 2). When comparing across sex and age the adducts that were identified through the Stanstrup database were carbamoyl-ethyl-dC, hydroxymethyl-methyl-Thy, oxohexenal-methyl-Ado, 7-pyridyl-hydroxy-butyl-dG, *cis*-butene-dial-dC, oxo-Cyd, as well as the canonical nucleobase Cyt, all adjusted to SG (Table 2). Putative chemical formulae were proposed for another 19 adducts that could not be identified through the Standstrup et al., database.^71^ Fragmentation patterns from each adduct were associated with sugar loss, indicating a ribose or 2-deoxyribose fragment, or a specific nucleobase. These characteristic fragmentation patterns were considered when determining the proposed chemical formula or identity. Each chemical formula proposed was within 5 ppm of the measured accurate mass. There remaining five adducts could not be identified nor could a chemical formula be proposed within 5 ppm of the measures m/z based on the characteristic fragments.

## Discussion

Urinary nucleic acid (DNA and RNA) adducts provide a noninvasive, potentially comprehensive assessment of the nucleic acid monomeric adduct burden and reflects the combined effects of adduct formation, DNA repair and systemic excretion.^72^ Experimental studies suggest that temporal patterns in adduct levels in urine may by influenced by the circadian regulation of DNA repair^73,74^ and fluctuations in renal clearance.^40,75–78^ For example, some previous studies directly examined circadian variation in urinary nucleic acid adducts ^31,79^ performed in two breast cancer survivors^59^ and six healthy individuals^80^, and used targeted analysis to determine the circadian variation of the adducts found in the urinary adductome^81^. These studies concluded that, for the adducts studied, first void spot urine sampling correlates with the gold standard 24-hour collection, and therefore spot urine samples could be used for biomonitoring urinary 8-oxodG. To our knowledge, the present study is the first to systematically evaluate diurnal fluctuations in urinary nucleic acid adduct excretion in a human population using an untargeted adductomics approach.

Our findings demonstrated that although global urinary adductome composition in number of adducts did not vary significantly with diurnal cycling, three of the 130 adducts exhibited higher peak intensities in first-void samples than in afternoon samples. These patterns were consistent across normalization strategies, indicating that the observed variation is not an artifact of data processing but reflects underlying biological processes.

### Intra- and inter-individual variability in the urinary concentration of representative nucleic acid adducts

The variability of any urinary nucleic acid adducts has rarely been systematically evaluated. To our knowledge, the studies that have evaluated diurnal variation have been highly targeted, focusing on 8-oxodG, 8-oxoGuo and/or 8-oxoGua.^40,75–78^ This predominance of inter-individual variability indicates that between-person differences exceed day-to-day fluctuations, supporting the use of these urinary adducts as exposure biomarkers, while also highlighting the contribution of within-person variability to overall measurement uncertainty.

To use an adduct as a reliable biomarker in population-based studies, the variability over time within an individual should be limited. As mentioned above, systematic diurnal fluctuations are limited. But preferably, adduct concentrations in urine should not vary much from day to day in the same individual, and any variability should be lower than the variability between individuals, resulting in a high ICC (0.8 or higher).^82^ In contrast, a low ICC limits the usefulness of a single urine sample, and instead repeated measurements are needed in epidemiological studies. Among the adducts evaluated within this study, 8-oxodG adjusted for creatinine or SG demonstrated acceptable ICC values exceeding 0.50. The adducts 7-Me-Gua and 2-Me-Cyt had low ICCs, indicating high intra-individual variability. Therefore, multiple samples would be required to generate reliable long-term estimates for their use in epidemiological studies.

The direction of diurnal trends differed between untargeted and targeted analyses: Although untargeted analysis indicated that most adduct intensities were generally unaffected by diurnal variation, three adducts showed significantly higher intensities in first void samples. In contrast, targeted analysis identified a few adducts which tended to have higher concentrations at afternoon sampling. The urinary concentrations of the three adducts selected for detailed analysis (8-oxodG, 7-Me-Gua, 2-Me-Cyt) were in agreement with previously reported ranges.^65–68^ Overall, our results suggest that diurnal variation, and thus sampling times, should be taken into consideration in epidemiologic applications of urinary nucleic adductomics.

### Untargeted variability in the urinary nucleic acid adductome

Multivariate analysis further supported our evidence of little diurnal variation. OPLS-DA models showed substantial overlap between first void and afternoon collected samples across adducts, with limited separation along the predictive component and dominant variance captured by orthogonal components. This pattern indicated that other sources of variability exceeded diurnal variation, at the level of the nucleic acid adductome. Collectively, these results indicate that diurnal regulation does not substantially alter the composition of the urinary nucleic acid adductome but instead modulates the relative abundance of a selection of adducts. In those cases it may be a consequence of temporal dynamic between adduct formation and clearance, which in turn is governed by a combination of diurnal fluctuations in metabolism,^74^ DNA repair,^73^ diuresis and renal function^83^, rather than diurnal fluctuations alone.

Both creatinine and SG are commonly used normalization approaches for urinary analyses, but the validity of these methods has not been systematically investigated for the totality of urinary DNA and RNA adducts. We found that neither method produced significant differences in the nucleic acid adductome. As previously reported, creatinine excretion increases with urinary flow rate,^39^ and flow rate itself is typically elevated in mid-day collections relative to first-void samples.^70^ Consequently, adjustment for creatinine could artificially reduce afternoon sample intensities, as the denominator of the adduct/creatinine ratio may increase disproportionately due to the impact of urine flow rate. Another limitation with adjustment for creatinine is the fact that creatinine excretion is affected by meat consumption and muscle mass.^84^ On average urinary creatinine concentrations are lower in women than in men (due to lower muscle mass)^84^, resulting in higher adduct/creatinine ratios in women even at the same excretion rates of adducts. In contrast, SG does not suffer from such potential problems. It is important to select an appropriate adjustment factor for diuresis when quantifying urinary biomarkers, particularly in time-resolved studies where physiological variables such as hydration,^85^ renal clearance,^83^ and metabolic rate^86^ may fluctuate. We did not identify any clear advantage of adjustment for creatinine or SG in the present data set, and therefore we adjusted for specific gravity for trends in adduct excretion.

Our findings further indicated that diurnal variation does not lead to changes in the number of adducts recovered in urine. Across all three sampling time points, the number of detected adducts remained consistent, suggesting that although diurnal variation may significantly impact the concentration of a few adducts, there is limited influence on the total adductome. Therefore, if the objective was to assess qualitatively the presence of specific adducts, a spot urine collected at any time of day would be sufficient. However, when evaluating quantitative differences, distinct temporal patterns may emerge depending on individual adduct identity, indicating that timing of collection should remain consistent across a study. We noted that the intensity of the majority of individual adducts was usually higher in the more concentrated first-void samples compared to the afternoon collection. This effect decreased when adduct intensities were adjusted for creatinine or SG when compared to non-normalized data, with only a limited number of adducts having higher intensities in the first void sample.

### Effect of sex and age on urinary nucleic acid adductome

Comparison of the adductome between age groups revealed divergent trends in males and females. In males, adduct concentrations were generally higher in the 21–30-y cohort, whereas in females, higher levels were observed in the 30–64-y population. This difference may arise from both hormonal and physiological factors. In men, maximum urine flow rates are reduced in younger adults during morning and evening hours compared to older individuals,^87^ potentially leading to longer urinary residence times and greater accumulation of repair-derived adducts in the bladder prior to excretion. In females, the opposing trend likely reflects endocrine changes associated with menopause. Perimenopause often begins in the fourth decade of life and transitions to menopause in the fifth, during which ovarian hormone production declines sharply.^88^ Estrogen influences both xenobiotic metabolism and renal function, modulating oxidative stress responses^89^ and urinary excretion patterns^53^; its loss may therefore alter the steady-state balance of adduct formation and clearance. A small subset of adducts remained more abundant in the younger female group, suggesting that these specific lesions may be less affected by hormonal regulation and follow patterns similar to those observed in males. Collectively, these data imply that age-related endocrine changes contribute to inter-individual variability in urinary nucleic acid adduct concentrations, particularly among peri- or post-menopausal women. It must be noted that the sizes of the sex- and age-stratified subgroups are small (n=10) and therefore these results should be interpreted with caution.

### Putative identification of variable adducts

Adducts that significantly varied over one or some of the comparisons made herein were further analyzed to determine putative identity. While the majority of adducts reported in this study remain unidentified, a number of adducts putatively identified are known to be byproducts of endogenous processes.^7,90^ These endogenously sourced adducts may also be exacerbated by exposure to one or some of the following: processed foods, cosmetics, vehicle exhaust, wood or coal burning stoves, and tobacco products, based on the putative identification of multiple adducts associated with these exposures. These variations support the observation that endogenously generated adducts are influenced by sex and age.

All the analyses presented were conducted at the population level, where data from individual adductomes were pooled to generate composite group profiles. The population observed here was part of a well-characterized biobank that has been used in previous biomarker studies,^39,56^ the participants were in generally good health and non-smokers. A limitation of the study was the small collection batch which only included three time points per individual, in addition to the modest sample size. Further studies incorporating larger and longitudinal cohorts are needed to confirm the patterns observed within sex and age groups, as well as to determine the nucleic acid adductome profiles associated with exogenous exposures, such as the ones listed above.

## Conclusions

Collectively, these findings emphasize that urinary nucleic adduct excretion is shaped by biological factors such as sex and age. Overall, diurnal variation was demonstrated by the increased adduct levels in samples collected at first void. For this reason, we recommend prioritizing first void urine collections given that this results from a long period of urine excretion and provides adducts at their highest concentration. First void urine will improve comparability across studies and minimize diurnal bias in future molecular epidemiological studies. Additionally, we found significant differences in adduct intensity and identity between males and females and established that there are significant differences in the intensities of the nucleic acid adductome between younger and older cohorts within the sexes; we suggest that this should be considered when analyzing samples from cohorts comprising of both males and females. Overall, this work demonstrates the potential for urinary nucleic acid adductomics to study diverse exposures in human populations across the age range, making it an ideal tool to dissect the exposome.

## Acknowledgements

The data presented in this publication were generated using instrument Orbitrap IQ-X Mass Spectrometer System with Vanquish Flex UHPLC System, which was supported by the National Institutes of Health (NIH) under Award Number S10OD036305, and a USF Interdisciplinary Research Award.

M.S.C., C-W.H and M-R.C. were supported, in part, by the National Institute of Environmental Health Sciences of the National Institutes of Health under grant number R01ES030557.

M.S.C. was supported, in part, by the National Institute of Environmental Health Sciences of the National Institutes of Health under grant numbers R01ES036931 and R44ES030274.

The content is solely the responsibility of the authors and does not necessarily represent the official views of the National Institutes of Health.

## CRediT

A.K., J.V., G.S and L.B: Writing – review & editing.

A.K. and M.M: Data curation, Visualization, Methodology, Investigation.

M.S.C., M-R.C., C-W.H: Conceptualization, Resources, Writing – review & editing, Supervision, Funding acquisition.

Y-J.C, L.S: Writing – review & editing

All authors reviewed and approved the manuscript.

