## Supplemental data for "Assessment of variability in the human urinary nucleic acid adductome"

**Table S.1. Isotopically labeled nucleic acid adducts were spiked into each sample as a reference cocktail of internal standards at a final concentration of 5 ppm.** Internal standards were prepared as individual solutions and as a reference cocktail with concentrations of 10 ppm in 1 mM ammonium acetate with 0.1 % formic acid.

| **Standard** | **Label** | **Structure** |
| --- | --- | --- |
| 5-methyl-2'-deoxycytidine | D_3_ | 2'-dN |
| 3’-O-methylthymidine | D_3_ | 2'-dN |
| 8-oxo-7,8-dihydro-2′-deoxyguanosine | ^13^C ^15^N | 2'-dN |
| 5-methylcytosine | ^13^C ^15^N_2_ | nB |
| 5,6-dihydrothymine | D_6_ | nB |
| 5-hydroxymethylcytosine | D_2_ ^13^C | nB |
| 3-methyladenine | D_3_ | nB |
| 9-methyladenine | D_3_ | nB |
| 6-O-methylguanine | D_3_ | nB |
| N7-(2-hydroxyethyl)guanine | D_4_ | nB |
| 9-(2-hydroxypropyl)adenine | D_6_ | nB |
| N3-(benzyl-D7)adenine | D_7_ | nB |
| 9-carboxymethoxymethylguanine | ^13^C_2_ ^15^N | nB |
| 2’-O-methylcytidine | D_3_ | rN |
| 2’-O-methyluridine | D_3_ | rN |
| 1-methyladenosine | D_3_ | rN |
| 2'-O-methyladenosine | D_3_ | rN |
| 2’-O-methylguanosine | D_3_ | rN |
| 2-(diethylamino)guanosine | D_6_ | rN |
| N6-succinyl adenosine | ^13^C_4_ | rN |

**Table S.2. Vanquish Horizon HPLC and Orbitrap IQ-X mass spectrometer system operating conditions.**

| Vanquish Horizon HPLC conditions | |
| --- | --- |
| Equipment | Thermo Scientific Vanquish Horizon HPLC system |
| HPLC column | Acclaim 120 C18 5um 120A 2.1 x 150mm |
| Column Temp | 50 °C |
| Aqueous | 1 mM ammonium Acetate with 0.1 % Formic Acid (mobile phase A) |
| Organic (mobile Phase B) | MeOH with 0.1 % Formic Acid |
| Gradient | 2% B (0 min – 2 min), 35% B (35 min), 50% B (55 min) – Equilibrium time 2% B (15 min) |
| Flow Rate | 0.100 mL/min |
| Injection Volume | 5 µL |
| IQ X mass spectrometer conditions | |
| Equipment | Thermo Scientific Orbitrap IQ X |
| Ionization Method | Positive mode ESI |
| Spray Voltage | 3.5 KV |
| Ion transfer tube temp | 275 °C |
| [Full MS] | |
| Resolution | 120 K |
| Mass Range | 100 – 600 m/z |
| [Data Dependent- MS^2^] | |
| Resolution | 30 K |
| Intensity Threshold | 10 k |
| Dynamic Exclusion | 6 s |
| Isolation Window | 1.2 |
| Automatic Gain Control | 50 K |
| Normalized collision energy | 20 %, 35 %, 60 % |
